# Transcellular chaperone signaling is an intercellular stress-response distinct from the HSF-1 mediated HSR

**DOI:** 10.1101/2022.03.17.484707

**Authors:** Jay Miles, Sarah Townend, William Smith, David R. Westhead, Patricija van Oosten-Hawle

## Abstract

Organismal proteostasis is maintained by intercellular signaling processes including cell nonautonomous stress responses such as transcellular chaperone signaling (TCS). When TCS is activated upon tissue-specific knockdown of *hsp-90* in the *C. elegans* intestine, heat-inducible *hsp-70* is induced in muscle cells at the permissive temperature resulting in increased heat stress resistance and lifespan extension. However, our understanding of the molecular mechanism and signaling factors mediating transcellular activation of *hsp-70* expression from one tissue to another is still in its infancy. Here we conducted a combinatorial approach using transcriptome RNA-Seq profiling and a forward genetic mutagenesis screen to elucidate how stress-signaling from the intestine to the muscle is regulated. We find that the TCS-mediated “gut-to-muscle” induction of *hsp-70* expression is suppressed by HSF-1 and instead relies on *transcellular-X-cross-tissue (txt)* genes. We identify a key role for the PDZ-domain guanylate cyclase *txt-1* and the homeobox transcription factor *ceh-58* as signaling hubs in the stress receiving muscle cells to initiate *hsp-70* expression and facilitate TCS-mediated heat stress resistance and lifespan extension. Our results provide a new view on cell-nonautonomous regulation of “inter-tissue” stress responses in an organism that highlight a key role for the gut. Our data suggest that the HSF-1-mediated HSR is switched off upon TCS activation, in favor of an intercellular stress-signaling-route to safeguard survival.

## Introduction

The preservation of protein homeostasis (proteostasis) is central for the maintenance of cellular and organismal health during environmental and physiological challenges. In multicellular organisms, intercellular signaling processes are essential for organismal development, differentiation and cell growth (Housden and Perrimon, 2014), as well as for the maintenance of organismal proteostasis (van Oosten-Hawle et al., 2013; Miles et al., 2019; Prahlad et al., 2008; Taylor and Dillin, 2013). The cell nonautonomous regulation of the heat shock response (HSR), the unfolded protein response of the endoplasmic reticulum (UPR^ER^) and the mitochondria (UPR^mito^) (Berendzen et al., 2016; Frakes et al., 2020; Prahlad et al., 2008; Taylor and Dillin, 2013) is playing a key role in the coordination of proteostasis across tissues.

The nervous system has a unique role in this process as it integrates neuronal stimuli for the transmission of a stress response to a distal tissue. It achieves this through wired synaptic connections whereby neurotransmitters such as serotonin or octopamine and tyramine function as key regulators of the cell nonautonomous HSR or the cell nonautonomous UPR^ER^, respectively (Bentley et al., 2016; Das et al., 2020; Imanikia et al., 2019; Özbey et al., 2020; Prahlad et al., 2008; Tatum et al., 2015; Volovik et al., 2014). Non-wired neuronal connections such as neuropeptides and Wnt signaling are also involved in the cell nonautonomous regulation of the UPR^ER^ and UPR^mito^ (Frakes et al., 2020; Zhang et al., 2018). In addition to the nervous system the *C. elegans* gut, being a major secretory organ, is another key tissue central for the regulation of organismal proteostasis. It achieves this via the release of neuropeptides, metabolites (O’Donnell et al., 2020; Shin et al., 2020; Zhang et al., 2013), innate immune peptides (Chikka et al., 2016; Gallotta et al., 2020; O’Brien et al., 2018; Peterson et al., 2019) as well as via lysosomal signaling (Imanikia et al., 2019). However, we do not know the identity of specific signaling cues activated by the gut that result in the upregulation of proteostasis regulators, such as chaperones in different tissues.

We have previously identified transcellular chaperone signaling (TCS) as a cell-nonautonomous stress response mechanism that mediates the activation of protective chaperone expression from one tissue to another (van Oosten-Hawle et al., 2013; O’Brien et al., 2018). TCS is induced by altering the expression levels of the molecular chaperone Hsp90 in specific tissues (van Oosten-Hawle et al., 2013). For example, neuron- or gut-specific overexpression of Hsp90 leads to a compensatory upregulation of the same chaperone in muscle cells (van Oosten-Hawle et al., 2013) which safeguards against chronic stresses such as age-associated amyloid protein misfolding (O’Brien et al., 2018). This is achieved via the activation of the transcription factor PQM-1 in the neurons and the intestine that upregulates extracellular innate immune peptides such as *clec-41* to coordinate organismal proteostasis via TCS (O’Brien et al., 2018). Conversely, reducing Hsp90 expression in the nervous system or the gut leads to the cell nonautonomous upregulation of *hsp-70* (*C. elegans* Hsp72/HSPA1A) that protects *C. elegans* from heat stress (van Oosten-Hawle et al., 2013).

Hsp90 is involved in the negative regulation of heat shock factor 1 (HSF-1) and the cytosolic heat shock response (HSR) (Anckar and Sistonen, 2011; Wu, 1995). Being part of a multichaperone complex, Hsp90 is involved in sequestering HSF-1 monomers in the absence of stress, and contributes to the deceleration of HSF-1 activity after a sufficient amount of heat shock proteins have been induced following stress (Anckar and Sistonen, 2011; Zou et al., 1998). Hsp90 inhibition leads to HSF-1 activation and results in the upregulation of heat shock proteins, including heat-inducible Hsp70 (Bagatell et al., 2000; Lindquist, 1986; Masser et al., 2020; Pincus, 2020). However, it is not clear whether this or a similar mechanism is induced upon tissue-specific Hsp90 knockdown that regulates transcellular activation of HSF-1 and *hsp-70* induction across tissues in *C. elegans*.

Here we examined how knockdown of *hsp-90* in the *C. elegans* intestine induces TCS-mediated expression of *hsp-70* in muscle cells. Using a combinatorial approach by analyzing gene expression profiles and a forward genetic screen, we identified *txt* genes and the homeodomain transcription factor CEH-58 as important mediators for TCS between the gut and the muscle. Surprisingly, TCS-mediated induction of *hsp-70* in muscle cells is suppressed by HSF-1 and requires the transcription factor CEH-58. Conversely, CEH-58 suppresses the HSF-1 mediated HSR. This antagonistic regulatory relationship between both transcription factors ensures only one type of organismal stress response is induced to mediate heat stress survival in *C. elegans*. Thus, our data shows that TCS is an organismal stress response distinct from the canonical HSF-1 mediated HSR that activates *hsp-70* expression in an HSF-1 independent manner.

## Results

### Intestine-specific knockdown of *hsp-90* induces TCS-mediated *hsp-70* expression and extends lifespan

We have previously shown that tissue-specific knockdown of *hsp-90* in the gut and neurons results in the cell-nonautonomous upregulation of *hsp-70* in multiple tissues of *C. elegans*, a process we termed Transcellular Chaperone Signaling (TCS) (van Oosten-Hawle et al., 2013). To further investigate how *hsp-70* expression is activated from one tissue to another, we used *C. elegans* strains expressing either an intestine-specific (*hsp-90*^*int*^*) or neuron-specific* (*hsp-90*^*neuro*^) hairpin RNAi construct against *hsp-90* (van Oosten-Hawle et al., 2013; Miles and Oosten-Hawle, 2020). Both strains show a particular induction of the heat-inducible *hsp-70p::mCherry* reporter in the bodywall muscle at the permissive temperature (20°C), corresponding to a 170-fold induction of *hsp-70p::mCherry* fluorescence intensity in the *hsp-90*^*int*^ strain and a 40-fold induction in the *hsp-90*^*neuro*^ strain **(Fig. 1A & 1B)**. The *hsp-70p::mCherry* reporter is heat-inducible, therefore no mCherry fluorescence is detected at 20°C in control animals that lack the *hp-RNAi* construct (*hsp-90*^*control*^) **(Fig. 1A)**. A 1-hour HS (35°C) induces the *hsp-70p::mCherry* reporter 4-fold in *hsp-90*^*control*^ animals **(Fig. 1B)**, primarily in spermatheca, intestine and pharynx as reported previously **(Fig. 1A)** (Guisbert et al., 2013), highlighting that the *hsp-70* tissue expression pattern induced by external HS is different from that induced by tissue-specific *hsp-90* knockdown. We confirmed induction of *hsp-70* expression and knockdown of *hsp-90* expression by measuring endogenous transcript levels using quantitative real time PCR. While knockdown of *hsp-90* in the intestine reduced whole-animal *hsp-90* mRNA levels by 50% **(Fig. 1D)**, endogenous *hsp-70* transcripts were induced 4-fold compared to *hsp-90*^*control*^ animals **(Fig. 1C; P<0.05)**. In the *hsp-90*^*neuro*^ strain *hsp-70* mRNA levels were induced 2.5-fold **(Fig 1C; P<0.05)**, although *hsp-90* transcripts were not measurably reduced, despite a detectable induction of *hsp-70* **(Fig. 1D)**. Overall, these results demonstrate that tissue-specific *hsp-90* hairpin RNAi leads to constitutive upregulation of *hsp-70* at the permissive temperature, particularly in bodywall muscle cells.

**Figure 1.**
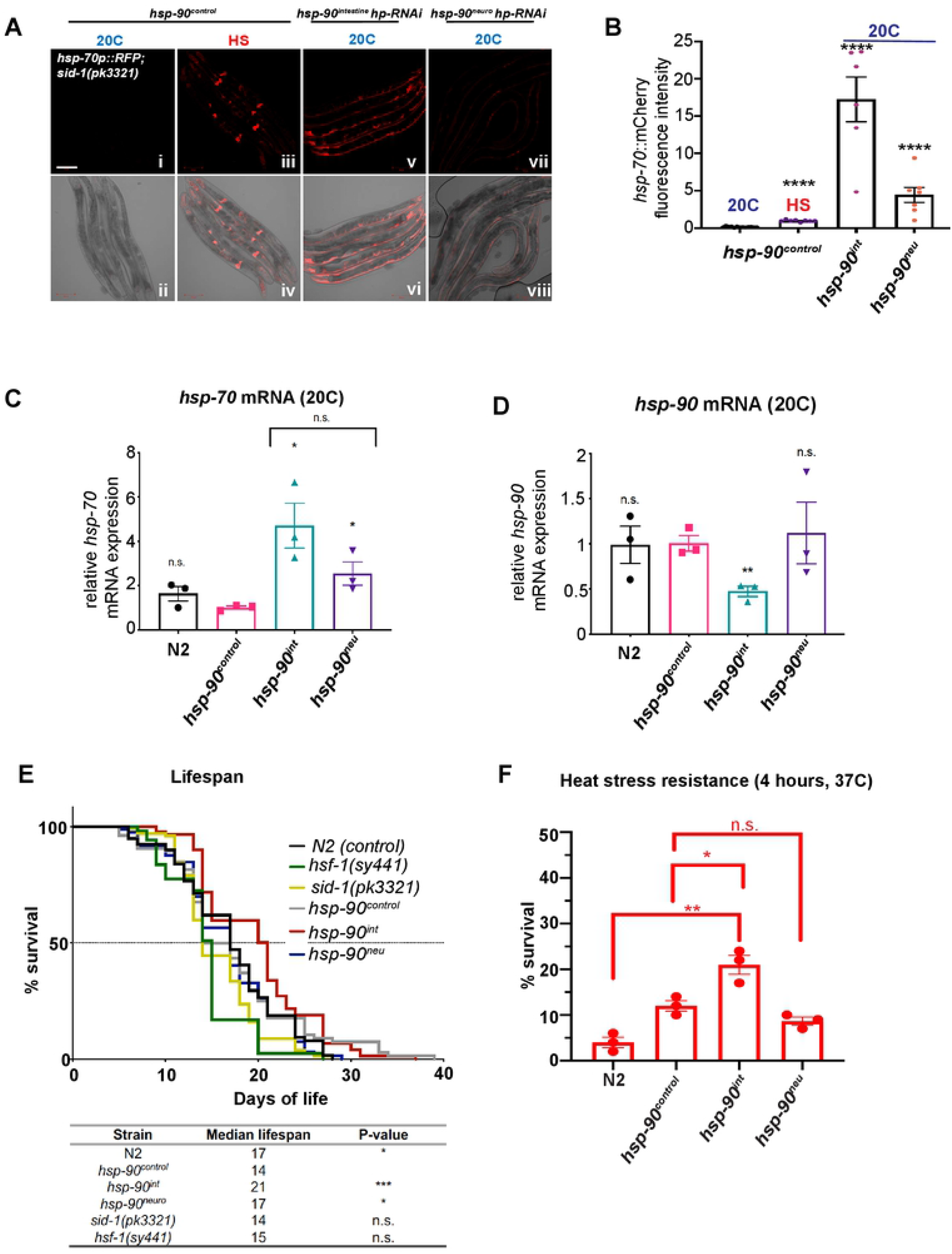
TCS induces *hsp-70* cell non-autonomously in muscle cells and increases lifespan and stress resistance. **A**. HS (1h; 35°C) induces expression of the *hsp-70p::mCherry* reporter in the spermatheca and pharynx (***iii, iv***). Intestine- (***v, vi***) and neuron-specific (***vii, viii***) *hsp-90* knockdown induces *hsp-70p::mCherry* expression in muscle cells at 20C. **B**. Quantification of *hsp-70p::mCherry* fluorescence intensity induced by TCS in *hsp-90*^*int*^ and *hsp-90*^*neu*^ compared to HS-induced *hsp-70p::mCherry* expression in control animals (*hsp-90*^*control*^). At least 5 animals per image and 3 biological replicates. Significance compared to mean fluorescence intensity in *hsp-90*^control^ was determined using student’s t test. **C**. Whole-animal *hsp-70* mRNA levels of *hsp-90*^*int*^ and *hsp-90*^*neu*^ animals and N2 compared to *hsp-90*^control^ nematodes at 20°C. **D**. Whole-animal *hsp-90* mRNA levels in *hsp-90*^*int*^ and *hsp-90*^*neu*^ animals compared to *hsp-90*^control^. **C & D**. Bar graphs represent the average of three biological replicates of 50 animals per RNAi and/or temperature condition. Error bars represent S.E.M of the 3 biological replicates. Significance compared to control *hsp-90*^control^ was determined using student’s t-test. **E**. Lifespan of *hsp-90*^*int*^ and *hsp-90*^*neu*^ compared to *hsp-90*^control^ and N2 animals. *hsf-1(sy441)* and *sid-1(pk3321)* strains were used as controls. n=100 animals per strain. Survival curves were compared using a log-rank (Mantel-Cox) test. **F**. Thermotolerance following a 2 and 4-hour heat shock at 37°C. n>3 replicates of 50 animals per strain per timepoint. Significance compared to *hsp-90*^control^ was determined using student’s t test. **B - F**.*P < 0.05; **P < 0.01; ***P < 0.001; ****P < 0.0001; n.s. = not significant.

To examine whether constitutive induction of *hsp-70* in *hsp-90*^*int*^ and *hsp-90*^*neuro*^ benefits *C. elegans* at the organismal level, we measured the effects on lifespan and resistance to heat stress. *hsp-90*^*int*^ animals showed a 50% increase in median lifespan (21 days; P>0.05) compared to *hsp-90*^*control*^ animals (14 days median lifespan; P<0.05), whereas no effect was measured in *hsp-90*^*neuro*^ animals **(Fig. 1E)**. By comparison, a *hsf-1(sy441)* hypomorph mutant that is deficient in its ability to induce heat shock proteins effectively shows a reduced median lifespan of 15 days, similar to *hsp-90*^*control*^ animals. This indicates that the *sid-1(pk3321)* mutation present in *hsp-90*^*control*^ affects lifespan could be suppressed by tissue-specific *hsp-90* knockdown **(Fig. 1E)**.

Exposure to heat stress (4h, 37°C) detrimentally reduced survival of wild type *C. elegans* (N2 Bristol) and *hsp-90*^*control*^ to 4% and 10%, respectively **(Fig. 1F)**. Interestingly, the thermotolerance of *hsp-90*^*int*^ animals was doubled (20%; P<0.05) compared to *hsp-90*^*control*^ animals, whereas *hsp-90*^*neuro*^ animals did not show an increased survival profile, perhaps mirroring the low level of TCS mediated *hsp-70* induction **(Fig. 1F)**. Overall, this showed that gut-to-muscle-mediated *hsp-70* induction was important to protect against the detrimental consequences of acute heat stress and leads to lifespan extension in *hsp-90*^*int*^.

### TCS-mediated *hsp-70* induction is suppressed by HSF-1

HSF-1 regulates the cytosolic heat shock response (HSR) and is required for the upregulation of heat-inducible *hsp-70* after HS or *hsp-90* knockdown (Anckar and Sistonen, 2011; Lindquist, 1986; Masser et al., 2020; Pincus, 2020; Zou et al., 1998). We therefore examined whether intestine- or neuron-specific *hsp-90 hp-RNAi* depended on functional HSF-1 to induce *hsp-70* in muscle tissue and increase organismal survival. To investigate this, we crossed *hsf-1(sy441)* mutants that cannot induce a proper HSR (Hajdu-Cronin et al., 2004) into the genetic background of *hsp-90*^*control*^, *hsp-90*^*int*^ and *hsp-90*^*neuro*^ animals expressing the heat-inducible *hsp-70p::mCherry* promoter. As expected, HS treatment (1h, 35°C) induced *hsp-70p::mCherry* fluorescence 10-fold in control (*hsp-90*^*control*^) animals (**Fig. 2D; P<0.0001**), and reduced it by 50% in *hsp-90*^*control*^*;hsf-1(sy441)* animals (**Fig. 2A & 2D; P<0.001**). Consistent with this observation, survival rates of *hsf-1(sy441)* and *hsp-90*^*control*^*;hsf-1(sy441)* animals decreased to 55% after a 2-hour exposure to 37°C and further dropped to below 10% after 4 hours of HS (**Fig. 2E**). Unexpectedly, the *hsf-1(sy441)* allele rendering HSF-1 dysfunctional, had no effect on *hsp-70* reporter expression in *hsp-90*^*int*^ and *hsp-90*^*neuro*^ animals at 20°C or after HS treatment, with *hsp-70p::mCherry* fluorescence intensity remaining constant during both conditions **(Fig. 2B, 2C and 2D)**. Moreover, *hsp-90*^*int*^*;hsf-1(sy441)* and *hsp-90*^*neuro*^*;hsf-1(sy441)* animals stably endured 2- and 4-hours of heat stress, with 26% (n.s.) of *hsp-90*^*int*^*;hsf-1(sy441) and 55%* (P < 0.01) *of hsp-90*^*neuro*^*;hsf-1(sy441)* animals surviving 4-hours at 37°C **(Fig. 2E)**. This result shows that TCS-mediated *hsp-70* upregulation enhances heat stress resistance and suggests that *hsf-1* acts as a suppressor of TCS. This indicates that TCS may be regulated by a molecular mechanism distinct from the canonical HSF-1 mediated HSR.

**Figure 2.**
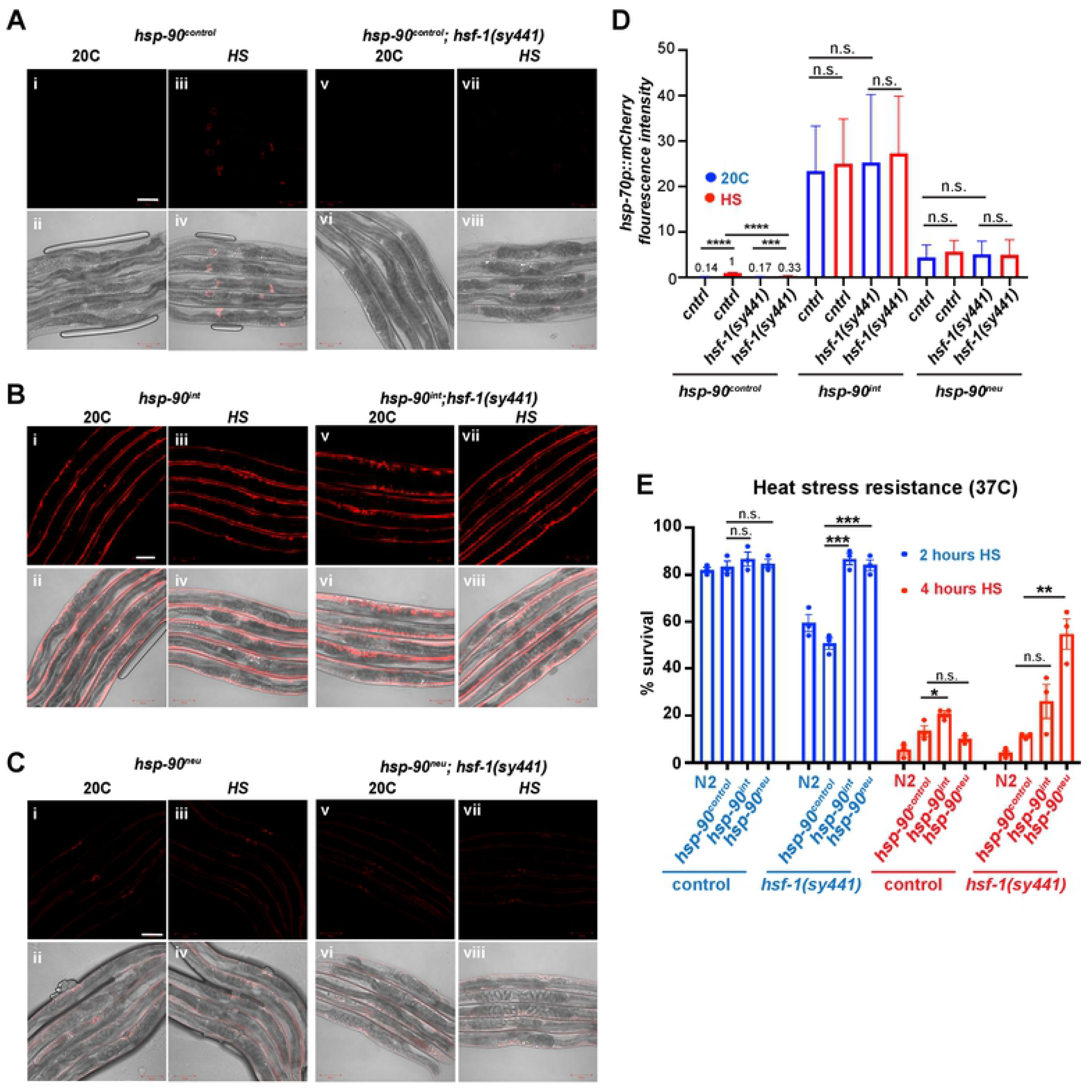
TCS-induced *hsp-70* expression is regulated independently of *hsf-1*. **A**. The *hsp-70p::mCherry* reporter is not expressed in *hsp-90*^*control*^ **(i, ii)** and *hsp-90*^*control*^*;hsf-1(sy441)* **(v, vi)** at 20°C. *hsp-70p::mCherry* is induced in *hsp-90*^*control*^ after HS **(iii, iv)**, but not in *hsp-90*^*control*^*;hsf-1(sy441)* mutants **(vii, viii). B**. Tissue-specific knockdown of *hsp-90* in the intestine (*hsp-90*^*int*^) induces the *hsp-70p::mCherry* reporter in the muscle at 20°C **(i, ii)** and after HS **(iii, iv)** to a comparable level. *hsp-70p::mCherry* expression in *hsp-90*^*int*^ animals before or during HS is independent of *hsf-1* (**v-viii**). **C**. Expression of the *hsp-70p::mCherry* reporter in *hsp-90*^*neuro*^ **(i, ii; v, vi)** and *hsp-90*^*int*^;*hsf-1(sy441)* animals **(iii, iv; vii, viii)** before and after HS. **D**. Quantification of *hsp-70p::mCherry* fluorescence intensity at 20°C or after HS in control (*hsp-90*^*control*^) or *hsp-90*^*int*^ and *hsp-90*^*neu*^ animals harboring wild type or mutated *hsf-1*. **E**. Survival of *hsp-90*^*int*^*;hsf-1(sy441)* and *hsp-90*^*neu*^;*hsf-1(sy441)* animals compared to *hsp-90*^control^ after a 2 and 4h HS at 37°C. N2 nematodes were used as an additional control. **D, E:** Bar graphs represent the average of three biological replicates of 50 animals per condition. Error bars represent S.E.M of the 3 biological replicates. Significance compared to *hsp-90*^*control*^ was determined using student’s t-test. * P<0.05; ** P<0.01; ***P<0.001; ****P<0.0001; n.s. = not significant.

### TCS is fundamentally different from the HSF-1 mediated HSR

To investigate how TCS differs from the HSF-1 mediated HSR, we first analyzed the transcriptional expression profile in *hsp-90*^*int*^ and *hsp-90*^*neuro*^ animals compared to the *hsp- 90*^*control*^ strain using RNA-Seq. In *hsp-90*^*int*^ worms, 281 genes were upregulated and 118 genes were downregulated at the permissive temperature (20°C) **(Fig. 3A)**, whereas *hsp-90*^*neuro*^ animals revealed a larger group of 1456 genes being upregulated and 795 genes downregulated **(Fig. 3B)**. The three most enriched GO-terms in *hsp-90*^*neuro*^ animals related to genes involved in neuropeptide signaling, the innate immune response, and transmembrane transport, suggesting a potential involvement in intercellular signaling processes **(Fig. 3D)**. *hsp-90*^*int*^ animals showed a clear enrichment for genes involved in the innate immune response and striated muscle contraction involved in embryonic body morphogenesis, which could be reflective of the strong *hsp-70p::mCherry* upregulation in the muscle in these animals **(Fig. 1A & Fig. 3C)**.

**Figure 3.**
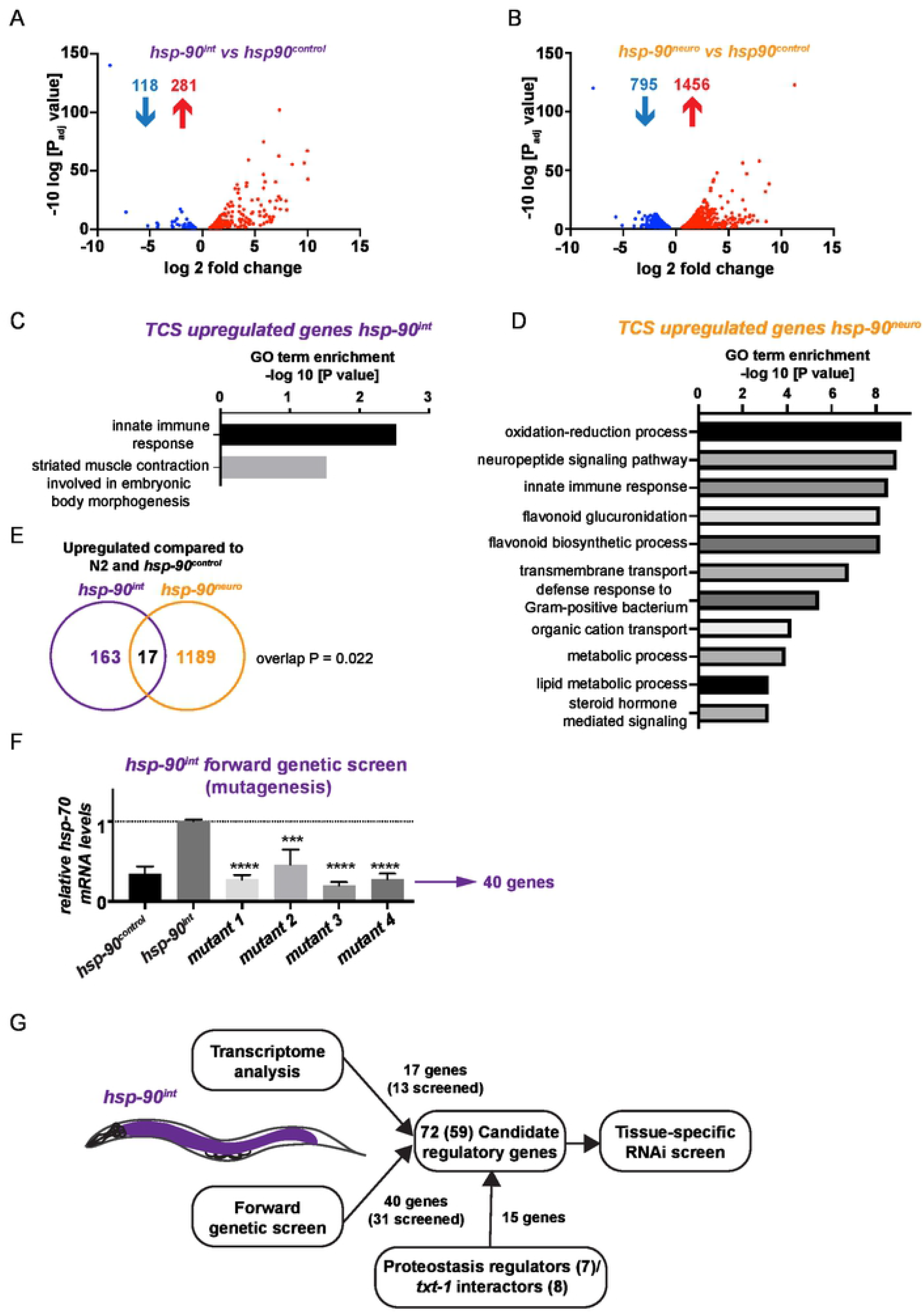
Transcriptional profiling (RNA-Seq) and a forward genetic mutagenesis screen identifies candidate genes required for TCS. **A**. RNA-Seq scatterplot showing log 2-fold change in expression levels of genes (P < 0.05) differentially regulated (118 downregulated; 281 upregulated) in the TCS activated strain *hsp-90*^*int*^ compared to *hsp-90*^*control*^. **B**. RNA-Seq scatterplot showing log 2-fold change in expression levels of genes (P < 0.05) differentially regulated (795 downregulated; 1456 upregulated) in the TCS activated strain *hsp-90*^*neuro*^ compared to *hsp-90*^*control*^. **C**. Gene ontology (GO) enrichment analysis of genes upregulated by TCS in *hsp-90*^*int*^. GO categories are reported as the -log 10 transformation of the P-value. **D**. Gene ontology (GO) enrichment analysis of genes upregulated by TCS in *hsp-90*^*neuro*^. **E**. Venn diagram analysis of genes upregulated in both *hsp-90*^*int*^ and *hsp-90*^*neu*^. P-value overlap = 0.022 was calculated using probability mass function of overlap size based on hypergeometric distribution. **F**. *hsp-90*^*int*^ mutants isolated after an EMS mutagenesis screen showing reduced global *hsp-70* mRNA expression that are comparable to *hsp-90*^*control*^ control animals. Whole-genome sequencing analysis identified 40 mutated genes in the isolated *hsp-90*^*int*^ mutants. See also Supplemental Figure 2. Bar graphs represent the average of three biological replicates of > 50 animals per strain; error bars represent S.E.M of the 3 biological replicates. Significance compared to the original strain *hsp-90*^*int*^ (pre-mutagenesis) was determined using student’s t-test. ***P<0.001; ****P<0.0001. **G**. Flowchart of identification of the 77 candidate genes resulting from the RNA-Seq analysis (17 genes), the mutagenesis & WGS analysis (40 genes) and proteostasis regulators and *txt-1* interactors (15 genes) that were taken further for a tissue-specific RNAi screen. Note that only 13 (out of 17) candidates of the RNA-Seq analysis and 31 candidates (out of 40) of the WGS analysis could be used for further RNAi analysis, as some candidate genes were not present in the genome-wide Ahringer RNAi library.

Both “TCS-activated” *hsp-90*^*int*^ and *hsp-90*^*neuro*^ strains **(Fig. 3E; Supp. Table 1)** share a set of 17 upregulated genes enriched for cell membrane proteins and extracellular soluble proteins, thus indicating a role for intercellular signaling processes that could regulate TCS **(Supp. Table 1)**. Interestingly, only few chaperones of the Hsp70 family were upregulated in *hsp-90*^*int*^ animals (*F44E5*.*4, F44E5*.*5* and *hsp-70*), while no chaperone genes were downregulated **(Supp. Fig. 1)**. Likewise, few chaperones were induced in *hsp-90*^*neuro*^ animals including small heat shock proteins (*hsp-12*.*3* and *hsp-12*.*6*) and four cadmium responsive ER chaperones (*cdr-2, cdr-4, cdr-5* and *cdr-6*) **(Supp. Fig.1)**. By comparison, none of the TCS-upregulated gene datasets were identified during HS conditions. Following HS, the most enriched GO-terms of HSF-1 upregulated genes are related to cuticle structure, translation and response to stress including members of the HSP16 (alphaB-crystallin) family of heat shock proteins (Brunquell et al., 2016). Comparison of the 815 genes induced by HS in wild type *C. elegans* (Brunquell et al., 2016) with our TCS dataset (*hsp-90*^*int*^) showed only a 1.7% overlap between the HSF-1 mediated HSR and TCS **(Supp. Fig. 1B; P < 0.04)**. Thus, the transcriptional program induced by tissue-specific knockdown of *hsp-90*, which activates TCS, is fundamentally different from the “canonical” HSF-1 mediated HSR that is triggered by external HS.

### Identification of candidate genes regulating TCS

In addition to RNA-Seq profiling, we undertook a forward genetic (mutagenesis) screen of *hsp-90*^*int*^ animals, to identify genes underlying the transcellular upregulation of *hsp-70* from the intestine to the muscle. We decided to focus on the effects of gut-to-muscle mediated TCS (*hsp-90*^*int*^ strain) for the mutagenesis screen, because of its strong *hsp-70p::mCherry* reporter expression in the bodywall muscle **(Fig. 1A**), which allowed for visual detection of increased or reduced mCherry fluorescence intensity. Four mutant strains were isolated showing reduced *hsp-70* reporter expression compared to the original *hsp-90*^*int*^ strain, indicating that these strains harbored a mutation in one or more genes required for TCS-mediated *hsp-70* upregulation **(Fig. 3F; Supp. Fig. 2A & 2B)**. To map the potential phenotype causing mutations, we performed whole-genome sequencing (WGS) combined with a SNP-based mapping step (Doitsidou et al., 2010) leading to the identification of 40 candidate genes (corresponding to 45 SNPs) that could potentially underlie reduced TCS (**Fig. 3F & Supp. Table 2**). Both WGS analysis and measurement of endogenous (whole-animal) *hsp-70* transcripts by qRT-PCR confirmed that the reduced *hsp-70p::mCherry* expression in these mutants was not due to a mutation in the promoter region of the *hsp-70*p::mCherry reporter (**Fig. 3F & Supp. Table 2**).

Together, the RNA-seq profiling identified 17 candidate genes and the forward genetic screen 40 candidates that could potentially regulate the TCS-mediated *hsp-70* activation from the intestine to the muscle. We named these candidate genes “*txt*” genes, “*<u>T</u>CS-cross(<u>X</u>)-<u>T</u>issue*” (**Fig. 3G; Supp. Table 1 & 2, 4)**.

### *txt* genes are required for TCS-mediated *hsp-70* induction from intestine-to-muscle

To determine in which tissue the *txt* candidate genes **(Supp. Tables 1 & 2)** act to facilitate TCS from the stress-perceiving “sender tissue” (intestine) to the *hsp-70p::mCherry* inducing “receiving tissue” (muscle) we performed a tissue-specific candidate RNAi screen (**see Fig. 3G and Fig. 4A for a flowchart**). In addition to the *txt* genes that were identified from the RNA-Seq and WGS analysis, we also included 15 genes known to be involved in the regulation of proteostasis including *pha-4, skn-1, pqm-1* and *daf-16* (van Oosten-Hawle et al., 2013; Hsu et al., 2003; Morley and Morimoto, 2004; O’Brien et al., 2018; Tullet et al., 2008), as well as previously identified interaction partners of *txt-1/C50D2*.*3* (8 genes) (Lenfant et al., 2010) as additional candidates for the tissue-specific RNAi screen **(Supp. Table 4)**. We utilized *hsp-90*^*int*^ strains that allowed for muscle-specific (strain PVH171) or intestine-specific (strain PVH172) RNAi-mediated knockdown of the 72 candidate genes and measured *hsp-70p::mCherry* fluorescence intensity as a read-out (Miles and Oosten-Hawle, 2020). Note that whole-animal RNAi cannot be used in *hsp-90*^*int*^ due to the *sid-1(pk3321)* mutant background that renders *C. elegans* insensitive to RNAi at the systemic level (Calixto et al., 2010; Miles and Oosten-Hawle, 2020; Winston et al., 2002).

**Figure 4.**
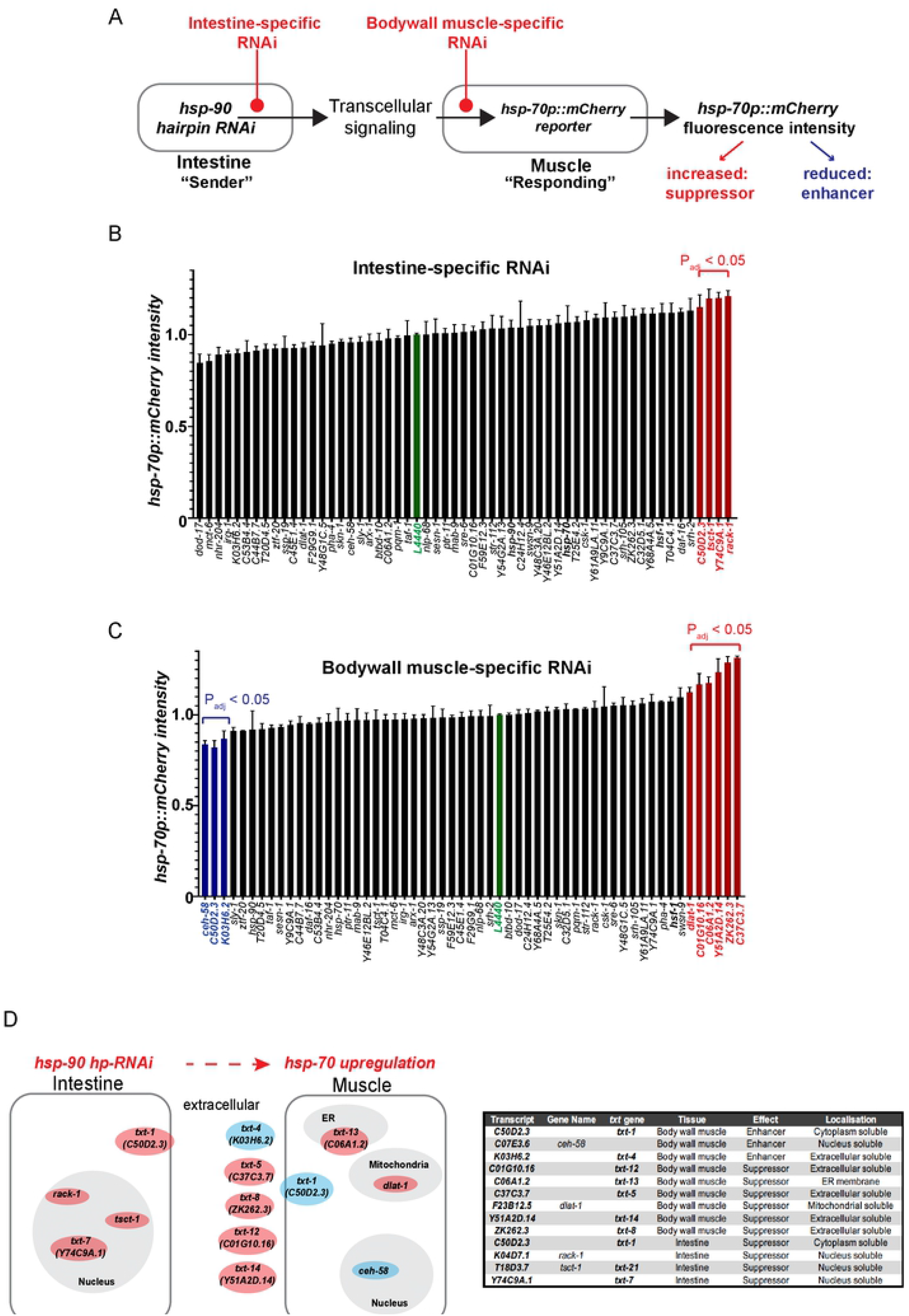
*txt* genes are required for TCS-mediated *hsp-70* induction from intestine-to-muscle. **A**. Schematic of the intestine- and muscle-specific RNAi screen using candidate genes that are either knocked down in the intestine, which expresses an *hsp-90 hp-RNAi* construct (sender tissue); or in the muscle where *hsp-70p::mCherry* fluorescence intensity is induced by TCS (“receiving tissue”). Differences in the level of TCS-induced *hsp-70p::mCherry* fluorescence intensity after candidate gene RNAi will point to regulators (enhancers or suppressors) of TCS. **B**. Intestine-specific candidate RNAi screen using *hsp-70p::mCherry* fluorescence intensity as a read-out. Red bars denote tissue-specific RNAi hits that result in a significantly increased fluorescence respectively (P_adj_; adjusted P value < 0.05). **C**. Muscle-specific RNAi screen using *hsp-70p::mCherry* fluorescence intensity as a read-out. Blue and red bars denote tissue-specific RNAi causing significantly reduced or increased fluorescence respectively (adjusted p value <0.05). **B &C**. Bar graphs represent the average of three biological replicates of 50 animals per condition. Error bars represent S.E.M of the 3 biological replicates. Significance compared to control RNAi (L4440, green column) determined by one-way ANOVA with correction for multiple testing using the two-stage linear step-up procedure of Benjamini, Krieger and Yekutieli. **D**. Diagram and table of predicted subcellular localizations of proteins encoded by *txt* genes, in the relevant tissues. Blue denotes fluorescence enhancers, red denotes suppressors. The five genes shown outside the body wall muscle encode predicted extracellular soluble proteins. *txt-1 (C50D2*.*3)* expressed in the cytosol, was identified as an intestine-specific suppressor and as a muscle-specific enhancer of TCS-mediated *hsp-70* induction in the muscle.

The intestine-specific RNAi screen identified four genes (*C50D2*.*3; tsct-1; Y74C9A*.*1* and *rack-1*) as intestine-specific suppressors of TCS-mediated *hsp-70p::mCherry* induction in the muscle **(Fig. 4B; P<0.05)**. The bodywall muscle specific RNAi screen identified three genes acting as muscle-specific suppressors (*C50D2*.*3; ceh-58; K03H6*.*2*) and six genes acting as enhancers (*dlat-1; C01G10*.*16; Y51A2D*.*14; ZK262*.*3; C37C3*.*7; C06A1*.*2*) of TCS-mediated *hsp-70* expression in the muscle **(Fig. 4C; P<0.05)**. Figure 4D summarizes the predicted subcellular localization of these gene hits (using the DeepLoc 1.0 web tool) and the tissue in which they act as enhancers or suppressors **(Fig. 4D)**.

Among the gene hits, *txt-1 (C50D2*.*3)*, raised specific interest as it appeared to function as a muscle-specific enhancer and an intestine-specific suppressor of TCS-mediated *hsp-70* expression **(Fig. 4B & 4C)**, suggesting a key role for *txt-1* in TCS. Interestingly, *txt-1* encodes for a predicted PDZ domain protein (Harris et al., 2010), which often act as scaffolds for larger multiprotein complexes at the inner cell membrane and are involved in transmembrane receptor organization and vesicle trafficking (Fanning and Anderson, 1996; Nourry et al., 2003). Such a function could be particularly relevant for intercellular signaling with *txt-1* acting as a key node at the plasma membrane of muscle cells that receives the intercellular signal to induce *hsp-70* expression. Moreover, the transcription factor *ceh-58*, a direct and known interactor of *txt-1* (Lenfant et al., 2010) was also identified as an enhancer of TCS-induced *hsp-70* expression in the muscle **(Fig. 4C)**, and encodes a homeobox transcription factor. Indeed, a CEH-58 consensus motif (TAATTA/G) is present in the promoter of the *hsp-70* gene 800 base pairs upstream of the *hsp-70* transcription start site ((Narasimhan et al., 2015); CIS-BP database; **Supp. Fig. 3A**). This suggested that *txt-1* and *ceh-58* could indeed be components of an inter-tissue signaling cue that transmits a TCS signal from the intestine to induce *hsp-70* expression in the muscle.

### Activation of TCS acts as a “switch” that shifts control of *hsp-70* expression from HSF-1 to TXT-1/CEH-58

To further examine the role of *txt-1* and *ceh-58* in TCS, we investigated how intestine- and muscle-specific RNAi of either gene affected *hsp-70* induction at 20ºC and HS, and whether this reduced the increased thermotolerance of *hsp-90*^*int*^ animals. Knockdown of *txt-1* or *ceh-58* by RNAi in muscle cells of *hsp-90*^*int*^ not only reduced expression of the *hsp-70p::mCherry* in the bodywall muscle as identified by the tissue-specific RNAi screen **(Fig. 4C)**, but also reduced global endogenous *hsp-70* transcripts at 20ºC and during HS **(Fig. 5A)**, as well as reducing thermotolerance by 50% **(Fig. 5D; P < 0.0001)**. This confirmed that both transcellular signaling components act as facilitators of TCS-induced *hsp-70* expression in muscle cells that is crucial for heat stress survival. Conversely, muscle-specific *hsf-1* RNAi increased *hsp-70* transcripts by 7-fold **(Fig. 5A)** and further improved HS survival rates to > 60% compared to a 20% survival rate of control animals **(Fig. 5D)**, verifying HSF-1 role’s as a suppressor of TCS. This suppressive function of *hsf-1* is however abolished when knocked down in combination with either *ceh-58* or *txt-1* by RNAi **(Fig. 5A)** suggesting that the interplay between *txt-1 & hsf-1* or *ceh-58 & hsf-1* is required for HSF-1’s repressive activity **(Fig. 5A)** as well as TCS-mediated survival rates **(Fig. 5D)**. Similarly, extracellular peptides *txt-4* and *txt-8* act as facilitators of TCS-mediated *hsp-70* induction in the muscle of *hsp-90*^*int*^ **(Supp. Fig. 4A)** and are required for thermotolerance **(Supp. Fig. 4D)**, indicating that both extracellular peptides could function as transmitters of TCS from the intestine to the muscle. In a control strain where TCS is not active and RNAi is systemic (strain AM722), *hsf-1* RNAi abolished *hsp-70* expression and thermotolerance as expected **(Fig. 5C and Fig. 5D)**. Strikingly, the opposite effect is achieved upon *ceh-58* RNAi in the same strain, resulting in a 10-fold induction of global *hsp-70* transcripts even at permissive temperature (**Fig. 5C)** and which also increased heat stress resistance **(Fig. 5D)**. Thus, while HSF-1 normally regulates the HSR by inducing *hsp-70* expression and survival in control animals exposed to HS, intestine-specific knockdown of *hsp-90* instead “switches off” the classic HSF-1 mediated HSR **(Fig. 5E)**.

**Figure 5.**
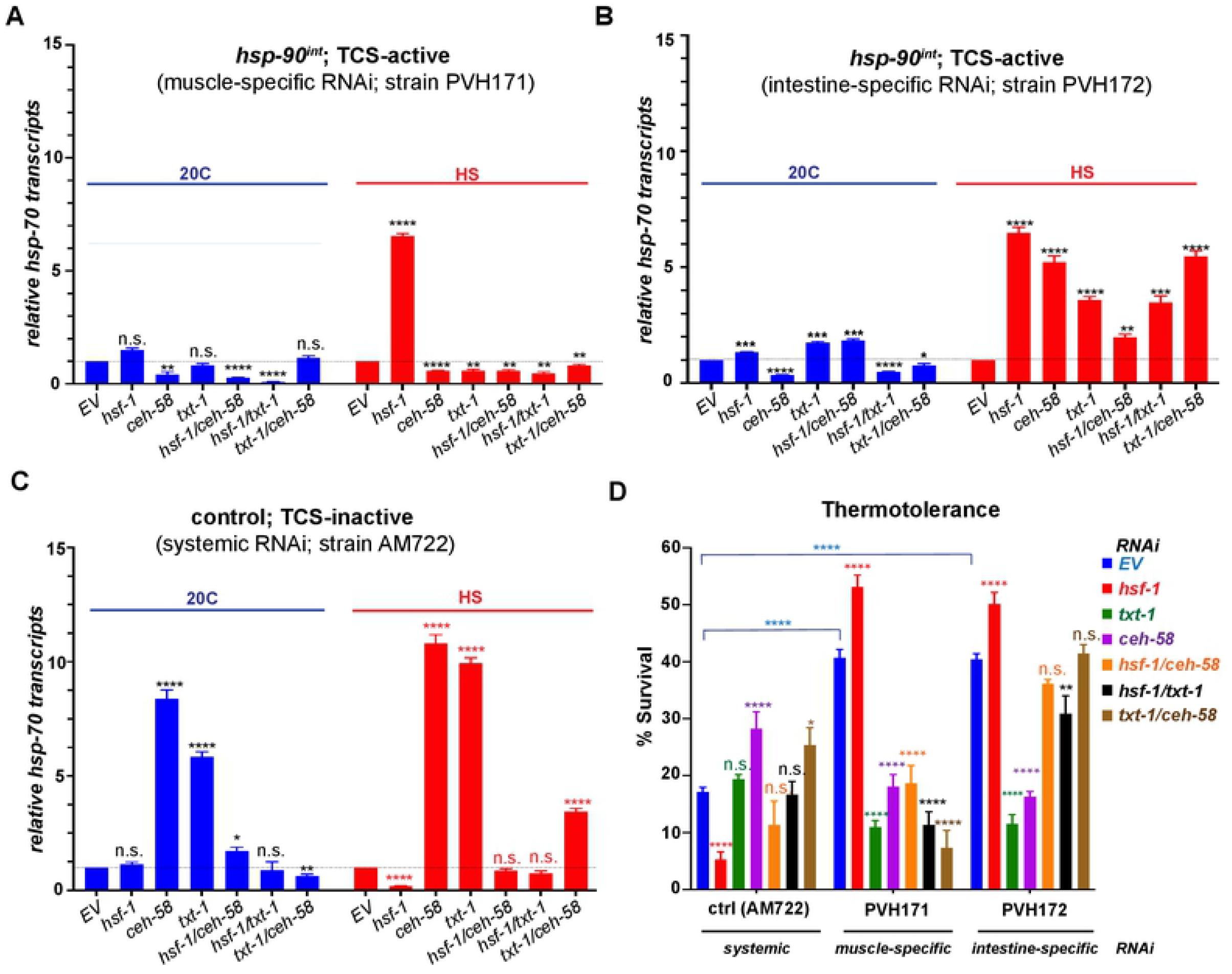
*txt-1* is required for TCS mediated *hsp-70* induction during HS independent of *hsf-1*. **A**. TCS-mediated *hsp-70* induction is facilitated by *txt-1* and *ceh-58* in the muscle. Quantification of whole-animal *hsp-70* transcripts in *hsp-90*^*int*^ animals at 20ºC and HS (35ºC), during muscle-specific RNAi in (strain PVH171) against *hsf-1, ceh-58, txt-1* and double RNAi of *hsf-1/ceh-58 hsf-1/txt-1, txt-1/ceh-58*. **B**. TCS-mediated *hsp-70* induction is suppressed by *txt-1* in the intestine. Quantification of whole-animal *hsp-70* transcripts in *hsp-90*^*int*^ animals at 20ºC and HS (35ºC), during intestine-specific RNAi (strain PVH172) against *hsf-1, ceh-58, txt-1* and double RNAi of *hsf-1/ceh-58 hsf-1/txt-1, txt-1/ceh-58*. **C**. *txt-1* and *ceh-58* are suppressors of the heat shock response. Quantification of whole-animal *hsp-70* transcripts in control animals at 20ºC and HS (35ºC), allowing for systemic RNAi (strain AM722) against *hsf-1, ceh-58, txt-1* and double RNAi of *hsf-1/ceh-58 hsf-1/txt-1, txt-1/ceh-58*. **D**. Thermotolerance after 6-hours of heat stress at 35ºC of TCS active strains during muscle-specific RNAi (strains PVH171) and intestine specific RNAi (strain PVH172) compared to a control strain (TCS-inactive; AM722) during systemic RNAi against *hsf-1, ceh-58, txt-1* and double RNAi of *hsf-1/ceh-58 hsf-1/txt-1, txt-1/ceh-58*. **A-D**. Bar graphs represent the average of three biological replicates of 50 animals per RNAi and/or temperature condition. Error bars represent S.E.M of the 3 biological replicates. Significance compared to control RNAi (EV) was determined using one-way ANOVA. * P<0.05; ** P<0.01; ***P<0.001; ****P<0.0001; n.s. = not significant.

The suppressive function of HSF-1 for TCS is also demonstrated in the intestine, as gut-specific *hsf-1* RNAi increased *hsp-70* expression at 20ºC and during HS **(Fig. 5B)**, as well as further enhancing thermotolerance **(Fig. 5D)**. The function of *txt-1* and *ceh-58* in the intestine however differs from their role in the muscle: in the intestine *txt-1* and *ceh-58* both suppress TCS-mediated *hsp-70* expression during HS, albeit *ceh-58* being a facilitator of *hsp-70* expression at 20ºC **(Fig. 5B)**. Interestingly, heat stress survival rates upon *ceh-58* and *txt-1* RNAi in the intestine are reduced **(Fig. 5D)**, despite increased global induction of *hsp-70* transcripts **(Fig. 5B)**. Intestine-specific knockdown of extracellular peptides *txt-8* and *txt-12* show a similar picture with reduced survival **(Supp. Fig. 4B)** despite increased TCS-mediated *hsp-70* expression **(Supp. Fig. 4E)**. This suggests that the inhibitory function of *txt-1/ceh-58* and extracellular peptides *txt-8* and *txt-12* in the intestine is required for TCS mediated regulation of survival.

We summarize these findings in a model describing gut-to-muscle stress signaling upon intestine-specific *hsp-90* knockdown and the consequences for heat stress survival **(Fig. 6)**. In control strains (“TCS-inactive”), *hsp-70* expression and heat stress survival are regulated in an HSF-1 dependent manner, with extracellular peptides TXT-4 and TXT-12 involved in this process. During HS, both TXT-1 and CEH-58 function as suppressors of HSF-1 mediated *hsp-70* induction **(Fig. 6B)**.

**Figure 6.**
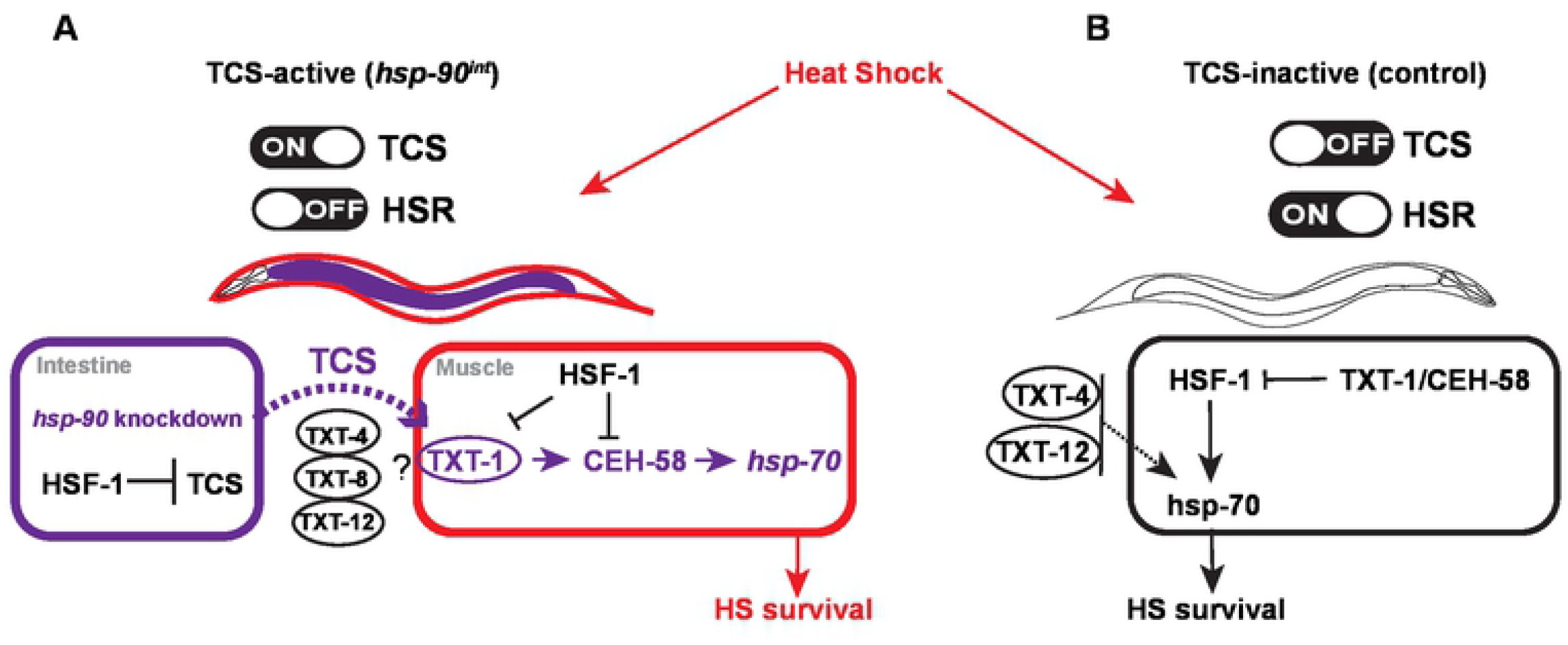
Proposed model explaining the opposing regulatory mechanisms of *hsp-70* induction and survival in TCS-active compared to a TCS-inactive strain. **A**. Upon *hsp-90* RNAi in the intestine (“TCS-active”) *hsp-70* is induced in the muscle via an *hsf-1*-independent mechanism that depends on TXT-1/CEH-58 signaling in the muscle as well as extracellular peptides TXT-4, TXT-8 and TXT-12. HSF-1 functions as a suppressor of TCS-mediated *hsp-70* expression and survival. **B**. In control animals (“TCS-inactive”), cell nonautonomous *hsp-70* induction and survival depends on HSF-1 and extracellular peptides TXT-4 and TXT-12 during HS; and is suppressed by TXT-1/CEH-58.

Knockdown of *hsp-90* in the intestine (“TCS-active strain”) relays a transcellular signal to muscle cells via extracellular peptides TXT-4, TXT-8 and TXT-12 **(Fig. 6A)**. In the muscle, the membrane-associated TXT-1 protein then facilitates intracellular receipt of the TCS signal towards the transcription factor CEH-58, which promotes *hsp-70* induction and organismal HS survival. TXT-1 may also be required to transduce a signal to HSF-1, as combinatorial knockdown of *txt-1* and *hsf-1* abolishes the suppressive function of HSF-1.

We conclude that TCS and the HSF-1-mediated HSR are regulated in an antagonistic manner, with TCS inhibiting HSF-1 activity, whereas conversely, HSF-1 suppresses TCS-induced TXT-1/CEH-58 signaling **(Fig. 6)**.

## Discussion

*Hsp-90* knockdown in the *C. elegans* intestine transmits a signal to induce *hsp-70* in muscle cells, by utilizing a hitherto uncharacterized signaling pathway. In this work, we identified *txt* genes and the homeodomain transcription factor CEH-58 as TCS mediators involved in the regulation between gut-to-muscle signaling. We find that TCS-mediated *hsp-70* induction relies on CEH-58 in the muscle, rather than HSF-1 which, interestingly, acts as a suppressor of TCS. *Vice versa*, CEH-58 suppresses *hsp-70* induction and survival following heat stress in TCS-inactive control strains. We propose that TCS and the HSF-1 mediated HSR are regulated in an antagonistic manner that distinguishes between intracellular stress (gut-specific *hsp-90* RNAi) versus external stress (heat shock) to safeguard and ensure organismal heat stress survival.

Inter-tissue signaling can be triggered in the gut to communicate with other tissues, such as during pathogenic infection (Chikka et al., 2016; Ewbank and Pujol, 2016; Peterson et al., 2019), or during FOXO-to-FOXO signaling that has a life-span extending effect via intestinal released lipid signals (Zhang et al., 2013). This role of the intestine has however received less attention in the context of cellular stress responses and cell nonautonomous induction of molecular chaperones. Our study here investigated such inter-tissue signaling initiated by the gut that results in the upregulation of *hsp-70* in muscle cells.

Lifespan extending effects and heat stress resistance as a consequence of *hsp-90* inhibition has been previously reported in *C. elegans*, by using small molecule drugs such as Radicicol (Monorden) and Tanespimycin (Janssens et al., 2019). However, this beneficial organismal consequence is only achieved if induced after *C. elegans* development is completed (at the L4 larval stage). Earlier Hsp90 knockdown at the systemic level otherwise leads to developmental arrest and dauer formation (Janssens et al., 2019; Somogyvári et al., 2018). In contrast, in our study intestine-specific *hsp-90* RNAi is induced constitutively from embryonic stages of development onwards, without developmental defects. This highlights apparent differences between systemic and tissue-specific knockdown of *hsp-90* and indicates that an intestinal role of *hsp-90* may not be required for some stages of development.

It is surprising that *hsp-90* knockdown in the intestine does not require HSF-1 to induce *hsp-70* expression in the muscle. Knockdown or inhibition of Hsp90 activity is known to activate HSF-1 mediated expression of heat-inducible Hsp70 (Wu, 1995; Zou et al., 1998). HSF-1 has been reported to facilitate intercellular communication between cancer associated fibroblasts that is required to promote cancer cell growth (Mendillo et al., 2012; Scherz-Shouval et al., 2014). This is in contrast to our finding in *C. elegans*, where HSF-1 functions as a suppressor of intercellular *hsp-70* induction upon gut-specific *hsp-90* knockdown. This suggests that at an organismal level more complex layers of regulation may be required to achieve the appropriate response in the correct target tissue. Perhaps the choice of an alternative transcription factor such as CEH-58 is necessary to achieve a high-level of specificity tailored for a specific cell type. This indicates the requirement for a specific “tissue-code” of organismal proteostasis, that ascertains induction of a cell-type specific stress response in metazoans.

Transcription factors other than HSF-1 are known to be involved in the regulation of heat shock proteins and molecular chaperones, including FOXO/DAF-16, SKN-1/NRF2, PHA-4 and PQM-1 (Blackwell et al., 2015; Hsu et al., 2003; Lindquist, 1986; O’Brien et al., 2018; Shpigel et al., 2019; Tepper et al., 2013). Although CEH-58 has previously not been implicated in chaperone expression, the *hsp-70* promoter contains a CEH-58 consensus sequence and our data suggests that CEH-58 genetically interacts with HSF-1. Future studies will need to confirm direct binding of CEH-58 to DNA elements in the promoters of stress-responsive heat shock proteins and a potential co-regulation with HSF-1 by direct protein-protein interaction between both transcription factors.

TCS clearly distinguishes itself from an HSF-1 mediated stress response, not only because TCS relies on CEH-58 for heat stress resistance but importantly because the transcriptional profile induced by TCS has little in common with the classic HSR mediated by HSF-1 (Supp. Fig. 1B). When TCS is induced in the intestine, innate immune response genes are a major upregulated gene group, rather than HSF-1 dependent heat shock proteins that are amongst the highest upregulated genes following external heat stress in *C. elegans* (Brunquell et al., 2016). Moreover, TCS induction in the intestine appears to “switch off” the HSF-1 mediated HSR in favor of a TXT-1/CEH-58 mediated signaling route that promotes protective *hsp-70* induction and survival. It is possible that these opposing effects are part of a built-in negative regulatory mechanism that safeguards stress survival.

In addition to CEH-58, TXT-1 is another signaling component, newly identified in this study to respond to intestinal derived TCS signals in muscle cells. TXT-1 is a PDZ (PSD-95, Discs-large, ZO-1) domain containing protein, which serve as signaling scaffolds for efficient and specific signal transduction at defined subcellular sites (Nourry et al., 2003; Roh and Margolis, 2003). TXT-1 itself is predicted to be localized at the cell membrane, and is expressed in muscle cells and the nervous system in *C. elegans*, similar to CEH-58 (Cao et al., 2017). Our data shows that TXT-1 and CEH-58 genetically interact as part of an **intra-**cellular signaling cue in the *C. elegans* bodywall muscle, corroborating a previous yeast-two-hybrid study that demonstrated direct interaction between both proteins (Lenfant et al., 2010). Although our study is the first to note an association of TXT-1 with the heat stress response in *C. elegans*, its closest human orthologue, DLG5, is a membrane associated guanylate cyclase that is part of the Hippo pathway (Venugopal et al., 2020; Zhu et al., 2016) and is involved in the cellular response to heat stress (Luo et al., 2020).

Extracellular peptides may be important contributors of organismal proteostasis that enable intercellular signaling (Melo and Ruvkun, 2012). Among extracellular peptides identified in our study, TXT-4 is a predicted extracellular lipase expressed in the intestine (Cao et al., 2017) that is crucial for the response to heat stress even in the absence of TCS. While it is an enhancer of *hsp-70* in muscle cells, it acts as a suppressor in the intestine, perhaps as a negative feedback mechanism required for gut-to-muscle stress transmission. Another extracellular TCS mediator is TXT-8, a phospholipase implicated in autophagy, which was previously identified as one of the 57 extracellular proteostasis regulators in *C. elegans* (Gallotta et al., 2020). Given their potential role in intercellular stress signaling, it will be interesting to follow up on the involvement of these extracellular proteostasis regulators in the regulation of cell nonautonomous stress responses (Gallotta et al., 2020). Interestingly, lipid signals have been previously suggested in FOXO-to-FOXO signaling from the intestine to muscle tissue (Zhang et al., 2013), suggesting a potential wider role for lipases such as TXT-4 and TXT-8 in the regulation of intercellular stress signaling.

Overall, our study highlights that transcellular signaling between the gut and muscle induces an HSF-1-independent stress response in muscle cells. We suggest that the opposing effects between TCS and the HSF-1 mediated HSR are part of a built-in negative regulatory mechanism benefiting organismal survival by monitoring responses in the most effective way. Further understanding of signaling elements involved in global stress responses that can be initiated in the gut to other tissues will become important for manipulation to promote organismal health. While this study focused on signaling occurring from the gut to the muscle, similar responses might exist in gut-to-brain signaling that could have potential major implications for the treatment of neurodegenerative diseases.

## Acknowledgements

We thank members of the P.v.O.-H. lab for critical reading of manuscript. *C. elegans* strains were kindly provided by the *Caenorhabditis* Genetics Center, funded by the NIH Office of Research Infrastructure Programs (P40 OD010440). Strains AM722 and AM994 were kindly provided by the Morimoto lab. This work was funded by the NC3Rs to P.v.O.-H. (NC/P001203/1). J.M. and S.T. were supported by an MRC Discovery Medicine North (DiMen) doctoral training partnership (MR/N013840/1).

## Author contribution

J.M and P.v.O.-H. designed the study, J.M., W.S. and S.T. performed the experiments and analysed the data, J.M. and D.R.W. analysed the RNA-Seq and whole-genome sequencing data, J.M. and P.V.O.-H. wrote the paper.

## Declaration of Interest

The authors declare no competing interests.

## Material and Methods

### *C. elegans* maintenance and strains

*C. elegans* strains used in this study are listed in Supplemental Table 3. Worms were maintained at 20°C on NGM agar plates seeded with OP50-1 *E. coli* according to standard procedures, unless otherwise stated (Brenner, 1974).

### Generation of transgenic strains

Extrachromosomal arrays expressing the neuron- and intestine-specific *hsp-90* hairpin constructs in a *sid-1(pk3321)* mutant background and expressing the *hsp-70p::mCherry* reporter (strain AM994) (van Oosten-Hawle et al., 2013) were integrated and backcrossed 5 times with strain AM994, that allows visualization of *hsp-70* induction via the stress-inducible *hsp-70p::mCherry* reporter. This resulted in strains PVH1 and PVH2 (see Supplemental Table 3).

For generation of PVH5 and PVH65, intestine specific (*vha-6p::SID-1::unc-54 3’UTR*) and muscle specific (*myo-3p::SID-1::unc-54 3’UTR*) SID-1 constructs were microinjected into AM994 and integrated and backcrossed 5 times, before crossing into PVH2. The ability of tissue-specific SID-1 expression to allow tissue-specific knockdown by feeding RNAi bacteria was confirmed as described in (Miles and Oosten-Hawle, 2020; O’Brien et al., 2018).

To perform tissue-specific RNAi experiments in the *hsp-90*^int^ strain, which carries a *sid-1* (*pk3321*) mutation preventing transport of RNAi between tissues, *sid-1* was reintroduced under the control of a tissue-specific promoter to facilitate tissue-specific RNAi uptake. This was achieved by outcrossing the *hsp-90*^int^ (PVH2) strain into the PVH5 and PVH65 strains, which express integrated constructs of *sid-1* under the control of either the *myo-3* body wall muscle-specific promoter or the *vha-6* intestine-specific promoter respectively in a *sid-1 (pk3321)* background, resulting in strains PVH171 and PVH172 (see Supplemental Table 3).

### Confocal microscopy Imaging

An inverted Zeiss LSM880 laser scanning confocal microscope was used to image *C. elegans*. Five worms were immobilized on 2 % agarose pads using 5 mM levamisole and a coverslip. Images which were subsequently used to quantify fluorescence were taken using a 10x objective. Quantification of fluorescence was performed using ImageJ software as described in (Miles and Oosten-Hawle, 2020). To image strain AM722 following heat shock, Day 1 adults were then incubated at 35°C for 1 hour followed by 3 hours recovery at 20°C.

### RNA extraction, cDNA synthesis and quantitative PCR (qPCR)

Nematodes were collected from NGM-Agar plates using chilled RNase-free water and worms were washed 3 times with RNase-free water to remove any residual bacteria (3 min, 500 x g). Excess water was removed and the pellet frozen at -80°C. TRIzol was added to samples before homogenisation using a pellet grinder, following which RNA was extracted using a Direct-Zol RNA MiniPrep kit (Zymo Research, Cambridge Biosciences). RNA concentration was measured using a Thermo Scientific NanoDrop One, and 100 ng of RNA was reverse transcribed into cDNA using a Bio-Rad iScript cDNA synthesis kit. Quantitative PCR was performed using Bio-Rad Universal SYBR green Supermix in a Bio-Rad CFX Connect Real-Time System. Relative transcript expression for each gene was determined using the delta-delta C_t_ method as described previously (van Oosten-Hawle et al., 2013; O’Brien et al., 2018). 3 biological replicates were performed per sample. Significance was determined using Student’s t test or one-way ANOVA with a cutoff of p < 0.05.

### Primers used for q RT PCR

*cdc-42 forw* 5’ TGTCGGTAAAACTTGTCTCCTG 3’

*cdc-42 rev* 5’ ATCCTAATGTGTATGGCTCGC 3’

*hsf-1 forw* 5’ GGACACAAATGGGCTCAATG 3’

*hsf-1 rev* 5’ CGCAAAAGTCTATTTCCAGCAC 3’

*hsp-70 forw* 5’ CGGTATTTATCAAAATGGAAAGGTT 3’

*hsp-70 rev* 5’ TACGAGCGGCTTGATCTTTT 3’

*hsp-90 forw* 5’ GACCAGAAACCCAGACGATATC 3’

*hsp-90 rev* 5’ GAAGAGCACGGAATTCAAGTTG 3’

### Lifespan assay

Animals were synchronized by egg-laying, and 100 L4-stage animals were selected per strain and transferred onto five NGM plates with 20 animals per plate. Every other day each nematode was assessed for survival, or censorship and numbers were recorded. Animals were assessed as dead if they did not display movement when gently touched on the nose using a platinum wire, and no pharyngeal pumping could be observed. Animals were censored if they crawled up the edge of the plate and became desiccated, burrowed into the agar, displayed the “bagging” phenotype where eggs hatched internally, displayed an exploded vulva phenotype, or if they could not be found. Any dead or censored animals were recorded and alive animals were transferred onto a new NGM plate. Data was analyzed using OASIS 2 online software (Han et al., 2016) and GraphPad Prism. Assays were repeated twice (2 biological replicates) and changes in lifespan were considered statistically significant when *P < 0*.*05* after a Log-Rank test analysis.

### Thermotolerance assay

Strains were synchronized by egg-laying, with gravid adults allowed to lay eggs for 6 hours and then removed from plates. Eggs were allowed to hatch and develop to L4 stage, at which point 3 replicate plates of 50 L4 animals were picked per condition per time point. The next day, plates containing day 1 adults were incubated at 35°C for either 6 hours or at 37°C for 2 hours; following which they were moved to 20°C. Animals were left to recover for 16 hours at 20°C, then survival of animals was scored. Animals were scored as alive if movement or pharyngeal pumping was observed. The percentage of alive and dead animals were scored and mean survival rates were determined using Student’s t test. Three independent experiments were performed for each strain (n=50 worms) and error bars indicate S.E.M.

### Transcriptomic profiling via RNA-seq

RNA was extracted from samples as described above. Agarose gel electrophoresis using a 1% gel was performed for a visual determination of sample quality, and RNA integrity number (RIN) was determined by the University of Leeds Next Generation Sequencing Facility using an Agilent 2200 TapeStation. RNA-seq was performed by Novogene (Hong Kong) on an Illumina Hi-Seq PE150 platform. Calculation of log_2_(fold change), p values and corrected p values were performed by Novogene. WBCel235 was used as the reference genome for annotation. GO term analysis was performed using the publicly available online tool DAVID Bioinformatics Resources 6.8.

### Forward genetic screen using EMS mutagenesis, whole-genome sequencing, and a Hawaiian *C. elegans* SNP mapping approach

*hsp-90*^int^ animals were synchronized by bleaching, allowed to develop to L4 stage, and incubated in 100 mM EMS solution for 4 hours at 20°C. 1000 mutagenized adults (P_0_ generation) were allowed to lay eggs (F_1_ generation) on NGM plates overnight and removed the following day. F_1_ eggs were allowed to mature and also lay eggs (F_2_ generation), following which adult F_1_ animals were removed. F_2_ animals were grown to L4 stage and screened under a fluorescent microscope for the desired phenotype of visibly altered *hsp-70p::mCherry* reporter fluorescence. In total approximately 20,000 F_2_ genomes were screened. Individuals identified by this method were isolated onto 35 mm plates and allowed to self-fertilize, and progeny monitored to ensure homozygous phenotypes. This identified candidate mutant strains 1-4 showing reduced *hsp-70p::mCherry* reporter fluorescence.

To identify mutations in these candidate strains which were potentially causal for the phenotypes of visibly reduced *hsp-70p::mCherry* fluorescence (mutants 1-4), a *C. elegans* Hawaiian SNP mapping approach as described in Doitsidou *et al*. 2010 was used (Doitsidou et al., 2010). The candidate populations identified through phenotypic screening, as well as the control strains (N2, *hsp-90*^control^ and *hsp-90*^int^), were each outcrossed to the Hawaii/CB4856 alternative wild-type strain. At the F_2_ stage of each of these outcrosses, 50 F_2_ animals were isolated and allowed to self-fertilize, and the 50 heterozygous populations subsequently recombined into a single sample for each outcross. These samples were frozen as pellets, from which genomic DNA was subsequently extracted using a Gentra PureGene Tissue Kit. Whole-genome sequencing of genomic DNA was performed by Novogene (Hong Kong) Company Limited on an Illumina Hi-Seq 2500 platform. WBCel235 was used as the reference genome for annotation.

Comparisons of data from each strain identified almost 15,000 SNPs as mutations. To determine which mutations were potentially causal for the phenotypes of interest, we used publicly available ‘CloudMap’ workflows on the online data analysis platform Galaxy, which we followed according to the user guide available in the CloudMap data library. The “CloudMap Hawaiian Variant Mapping with WGS and Variant” workflow pipeline (Minevich et al., 2012) was used to calculate the ratio of Bristol-derived to Hawaii-derived alleles at each SNP in each strain, and the “CloudMap Variant Discovery Mapping” workflow was used to remove SNPs which also occurred in control strains. Analysis was performed using the WormBase version WS266 which uses the WBcel235 reference genome. SNPs identified in this manner were taken forward as potentially causal if they had a Bristol-derived to Hawaii-derived allele ratio of less than 0.25 (Supplemental Table 2). The gene transcripts affected by these potential causal SNPs were identified using annotation performed by Novogene. Transcripts were excluded if they were classed as intergenic or pseudogenes.

### Gene knockdown by RNAi

Populations were synchronised by egg-laying on HT115 *E. coli* transformed with appropriate RNAi vectors (J. Ahringer, University of Cambridge, Cambridge, UK) over two generations. Synchronised F_2_ generation eggs were allowed to develop and were used in experiments as Day 1 adults.

## Data and Software availability

The data discussed in this study have been deposited in NCBI’s Gene Expression Omnibus and are accessible through GEO Series accession number GSE197412. https://www.ncbi.nlm.nih.gov/geo/query/acc.cgi?acc=GSE197412

## Supplemental Information

**Supplemental Figure 1. The TCS-induced transcriptional program is distinct from the HSF-1 mediated heat shock response**.

**A**. Differential chaperone gene expression of *hsp-90*^*int*^ and *hsp-90*^*neuro*^ compared to *hsp-90*^*control*^ strain. Lists of differentially expressed chaperones in each strain were compared to known *C. elegans* chaperone genes (Brehme et al., 2014). log2 FC (fold-change) compared to the *hsp-90*^*control*^ strain. P_adj_ = Bonferroni corrected P-value.

**B**. Venn diagram showing the overlap of 18 genes (1.7%) that are commonly upregulated between a TCS-active strain (*hsp-90*^*int*^) at 20ºC (this study) compared to N2 Bristol during HS (Brunquell et al., 2016). P-value overlap < 0.04 was calculated using probability mass function of overlap size based on hypergeometric distribution.

**Supplemental Figure 2. *hsp-70p::mCherry* fluorescence intensity of *hsp-90***^***int***^ **mutant strains isolated by EMS mutagenesis**.

**A**. Confocal images of the four *hsp-90*^*int*^ mutant strains (mutant 1-4) showing reduced *hsp-70p::mCherry* expression in the bodywall muscle compared to the parent strain *hsp-90*^*int*^. Scale bar = 100 μm.

**B**. Quantification of *hsp-70p::mCherry* fluorescence intensity in the EMS mutagenesis generated *hsp-90*^*int*^ mutant strains. *P < 0.05; **P < 0.01; ****P < 0.0001. 3 biological replicates per image with five or more animals per replicate. Significance compared to mean fluorescence intensity in *hsp-90*^int^ was determined using student’s t test.

**Supplemental Figure 3. The *hsp-70* promoter contains a CEH-58 consensus sequence**. Motif scanning in the *hsp-70* promoter identifies a consensus motif for the homeobox transcription factor CEH-58 of NNTAATTRNN (CIS-BP database, (Narasimhan et al., 2015, 2015). The CEH-58 consensus motif is located 800 base pairs upstream of the first ATG (marked CEH-58, blue). Scanning also identified two canonical heat shock elements (HSEs, marked in red) of the form TTCNNGAA at 106 and 758 base pairs upstream of the ATG.

**Supplemental Figure 4. Extracellular peptides are involved in the regulation of the HSR and TCS**.

**A**. Predicted extracellular peptides *txt-4* and *txt-8* are required in the muscle for TCS-mediated *hsp-70* induction during HS. Quantification of whole-animal *hsp-70* transcripts during muscle-specific RNAi (strain PVH171) against *txt-4, txt-5, txt-8, txt-12 and txt-14* compared to *EV* at 20ºC and during HS (35ºC). **B**. Predicted extracellular peptides *txt-4*, txt-5, *txt-8, txt-12, and txt-14* are suppressors of TCS-mediated *hsp-70* induction in the intestine during HS. Quantification of whole-animal *hsp-70* transcripts during intestine-specific RNAi (strain PVH172) against *txt-4, txt-5, txt-8, txt-12 and txt-14* compared to *EV* at 20ºC and during HS (35ºC).

**C**. Systemic RNAi-mediated knockdown of *txt-4* and *txt-12* in a “TCS-inactive” control strain (AM722) reduces *hsp-70* expression at 35ºC to ∼ 25% compared to control RNAi (EV), whereas *txt-5* and *txt-8* RNAi reduce *hsp-70* levels to 75% compared to EV RNAi.

**D**. Thermotolerance of *hsp-90*^*int*^ during muscle-specific RNAi (strain PVH171) against *txt-4, txt-5, txt-8, txt-12* and *txt-14* compared to control RNAi (EV). Day 1 adults were exposed to a 6-hour heat shock at 35ºC. **E**. Thermotolerance of *hsp-90*^*int*^ during intestine-specific RNAi (strain PVH172) against *txt-4, txt-5, txt-8, txt-12* and *txt-14* compared to control RNAi (EV). **F**. Thermotolerance of a control strain (TCS-inactive; AM722) during systemic RNAi against *txt-4, txt-5, txt-8, txt-12* and *txt-14* compared to control RNAi (EV). **A-E**. Bar graphs represent the average of three biological replicates of 50 animals per RNAi and/or temperature condition. Error bars represent S.E.M of the 3 biological replicates. Significance compared to control RNAi (EV) was determined using one-way ANOVA. *P<0.05; ** P<0.01; ***P<0.001; ****P<0.0001; n.s. = not significant.

**Supplemental Table 1. Genes upregulated in both TCS activated strains, *hsp-90***^***int***^ **and *hsp-90***^***neuro***^, **compared to wild type (N2) and *hsp-90***^***control***^. The subcellular localization was predicted from the amino acid sequence using the DeepLoc 1.0 webtool (https://services.healthtech.dtu.dk/service.php?DeepLoc-1.0).

**Supplemental Table 2. 45 SNPs (corresponding to 40 genes) identified by the mutagenesis screen and whole-genome-sequencing analysis**. All phenotype specific SNPs occurring in mutants 1-4 were ranked according to their CB4856: N2 ratio, with the lowest ratio of 0.139 ranked first. A ratio of 0, indicating one specific causal SNP was not identified. Multiple SNPs were identified in genes *Y48G1C*.*5, csk-1, ssp-19, sop-3* (grey shaded). LG, Position: chromosome and position of SNP on the genome. Reference genome WS266 or a combination of WS266 and WS220 were used to identify SNPs.

**Supplemental Table 3**. *C. elegans* strains used in this study.

**Supplemental Table 4**. Candidate genes used for the tissue-specific RNAi screen (see Figure 4).

## Notes

### Competing Interest Statement

The authors have declared no competing interest.

